# A Proof-of-Concept Preparation of Lipid-Plasmid DNA Particles Using Novel Extrusion Based 3D Printing Technology, SMART

**DOI:** 10.1101/2023.08.26.554956

**Authors:** Jaidev Chakka, Mohammed Maniruzzaman

**Affiliations:** Pharmaceutical Engineering and 3D Printing (PharmE3D) Lab, Division of Molecular Pharmaceutics and Drug Delivery, College of Pharmacy, The University of Texas at Austin, Austin, TX 78712, USA; Department of Pharmaceutics and Drug Delivery, School of Pharmacy, University of Mississippi, University, MS 38677

**Keywords:** Lipid nanoparticles, plasmid DNA, GFP, SMART, Transfection, Transient expression

## Abstract

Gene therapy is a promising approach with delivery of messenger RNA, small interference RNA, and plasmid DNA to elicit a therapeutic action *in vitro* using cationic or ionizable lipid nanoparticles. In the present study, a novel extrusion based Sprayed Multi-Adsorbed droplet Reposing Technology (SMART) developed in-house was employed for preparation, characterization, and transfection abilities of the green fluorescence protein (GFP) plasmid DNA in cancer cells *in vitro*. The lipid composition (ionizable) and plasmid DNA (pDNA, GFP) were mixed in a 1:1 ratio using SMART technology at 1, 5, 8 & 10 N/P ratios. The particles were characterized to determine particle size (DLS), zeta potential and morphology using scanning electron microscopy (SEM). The particle size yielded for all N/P ratios is in the range of 100 nm to 200 nm. The *in vitro* transfection was carried out in MG63 cells showed optimal formulation N/P 8 with expression of GFP protein. The toxicity study through MTT assay showed N/P 8 with toxicity lower than other groups. The results showed that the processes developed using SMART technology are consistent and can be utilized for commercial applications.

## 1. Introduction

Plasmids are one of the key elements in gene cloning and Recombinant DNA technology^1^. The earlier discovery that naked plasmid encoding luciferase gene can be expressed as protein in the muscle tissue of live mice^2^ has opened immense possibilities for plasmid based therapeutics^3^ and vaccines^4^. The naked plasmids are prone to nucleases and self-hydrolysis inside targeted cells limiting the potential of plasmid DNA from expressing enough protein to reach a therapeutic window for disease treatment. Hence the encapsulation of plasmids inside a nanoparticle is the logical choice for treatment modalities. Nanoparticle delivery vehicles such as liposomes, polyethyleneimine / cationic polymers are widely employed to create non-viral gene delivery strategies. Cationic liposomes were employed to delivery plasmid DNA where the particle size dependent endocytosis was observed^5^. Polyethyleneimine, both branched and unbranched polymers were a popular choice of plasmid DNA non-viral gene delivery vehicles. The cationic polymer was used for delivery of fibroblast growth factor (pFGF)^6^, bone morphogenic protein (pBMP2)^7^, vascular endothelial growth factor (pVEGF)^8^ and plasma derived growth factor (pPDGF)^9^ genes *in vitro* and *in vivo*.

Lipid nanoparticles (LNPs) have emerged as a novel and promising delivery vehicle upon its immense success with COVID mRNA vaccines. Besides, lipid nanoparticles were also employed to deliver a variety of therapeutic agents such as mRNA^10^, *si*RNA^11^, microRNA^12^ and plasmidDNA^13^. LNPs are a type of vesicle that has a homogeneous lipid core. The types and composition of the lipid is essential for complete encapsulation of the payload and effective internalization by cells. Ionizable lipids are one of the key component of the recently developed lipids that initiate electrostatic binding to endosomal membranes and trigger cytosolic release of gene moiety from the lipid nanoparticle^14^. However, for most of the lipid nanoparticle formation, it needs cationic or ionizing, phospholipids, structural and circulation enhancing lipids. Cationic or ionizable lipids initiate the first steps of self-assembly through electrostatic interactions. The toxicity of the cationic lipids has prioritized to utilize ionic lipids ex: DLin-MC3-DMA^15^ which are pH dependent for better efficacy and toxicity^16^. Cholesterol is a naturally abundant source present in cellular membranes at 20-50% provides structural stability to the lipid nanoparticles^17^. Phospholipids such as di-stearoyl phosphatidyl choline (DSPC) help packaging of the lipids into a stable nanoparticle where they constitute around 10-20% of a typical lipid nanoparticle formulation^18^. Polyethylene glycol (PEG) based lipids are the circulation enhanced lipids that control the half-life and cellular uptake of the LNPs. These lipids undergo modifications such as hyaluronan^19^ or antibodies^20^ to convert otherwise non-targeted delivery vehicle into a targeted delivery system. The ideal composition of the lipids with the right fabrication method could facilitate the generally accepted size range of 20-200 nm to withstand the fluid flow of blood and lymph^21^.

Lipid nanoparticles were prepared by mixing lipid (ethanolic) and pDNA (aqueous) components in different ratios and different nitrogen (lipid) to phosphorous (pDNA) (N/P) ratios to achieve complete encapsulation. There are several ways to achieve the LNPs by thin-film hydration, nanoprecipitation, and microfluidic mixing where the last one is very popular for lab scale and commercial fabrication. However, new mixing technologies such as impingement jets mixing (IJM) technology was widely used and being used by several big pharma companies producing COVID vaccines. Though popular microfluidic and impingement jets mixing technologies are not portable. The microfluidic chip fabrication involves a lot of cost imposing equipment where the viable span of the microfluidic chip is limited. The IJM is best suited for large-scale fabrication at the moment. Considering the vacuum in the area of portability for both small scale and large-scale fabrication with diverse nanoparticle preparation capability in preparing the LNPs, we have invented a new Sprayed Multi-Adsorbed droplet Reposing Technology (SMART) (WO2023283320A2) preparation method where the pneumatic extrusion of the three-dimensional (3D) printing was explored. 3D extrusion is majorly employed to prepare 3D scaffolds for bioprinting. With our novel and *state-of-the-art* technology, the crucial and delicate biologics can be prepared with greater than 100% encapsulation efficiency and no loss or degradation of the payload. The unique portability and consistency of this technology was confirmed through preparation of polylactide-co-glycolide^22^ and Chitosan^23^ nanoparticles for delivery applications *in vitro*, by our group.

In the present study, the ionizable lipid DLin-MC3-DMA having two dilinoleyl chains was used^24^. We used SMART to prepare lipid-plasmid DNA (GFP) nanoparticles, characterized and evaluated the toxicity and gene expression characteristics in MG63 cancer cells *in vitro*.

## 2. Experimental

The following lipids 1,2-distearoyl-sn-glycero-3-phosphocholine (18:0, DSPC), Cholesterol (Chol), 1,2-dimyristoyl-rac-glycero-3-methoxypolyethylene glycol-2000 (DMG-PEG) were procured from AvantiPolar, USA. The ionizable lipid (6Z,9Z,28Z,31Z)-Heptatriaconta-6,9,28,31-tetraen-19-yl 4-(dimethylamino) butanoate (DLin-MC3-DMA) was purchased from VWR, USA. All materials were used without any modifications.

### 2.1 Preparation of plasmid

The GFP plasmid (pEGFP_mEGFP-GluA1c) was a gift from Haining Zhong (Addgene plasmid # 114136; http://n2t.net/addgene:114136; RRID:Addgene 114136). The GFP plasmid engineered DH5α *E. coli* was (Addgene, USA) received from the manufacturer was streaked into the luria agar plates already mixed with Kanamycin (100 μg/mL). The plates were incubated at 37 °C overnight. The developed single colonies were inoculated into 10 ml Luria broth and cultured at 37 °C, 100 rpm for 8 h. The inoculum was transferred to 200 mL fresh media with kanamycin in a 500 mL flask and cultured for 48h. The bacteria were processed using plasmid extraction and purification kit (HiSpeed Plasmid Maxi kit, Qiagen, USA). The plasmid was extracted by following the kit instructions. The final extracted plasmid was dispersed in water and quantified at 260 nm (Nanodrop, Thermo Scientific, USA).

### 2.2 Preparation of Lipid plasmid particles

The lipids DSPC:Chol:MC3:DMG-PEG of 10:48:40:2 were mixed from the stock solutions of 6 mg/mL. The plasmid of 10 μg/mL was dissolved in 10 mM citrate buffer (pH 4). The lipid and plasmids were mixed in a 1:1 ratio to get the LNP particles. The SMART was carried out in the pneumatic extrusion enabled three-dimensional printer (Biox, Cellink, USA) where 1 ml of the lipid composition was extruded at 200 KPa into 1 mL of plasmid solution. The mixture was taken again into the syringe and extruded to ensure uniform extrusion and particle formation. To ensure maximum encapsulation/complexation of pDNA with lipids, the components were mixed at different N/P ratios where the lipid component was fixed while varying the pDNA (GFP) amount.

### 2.3 Characterization

The LNP-pGFP were analyzed for particle size dynamic light scattering (DLS) method and zeta potential in Zetasizer (Malvern, USA). The stability of the particles was tested by gel retardation assay. A 1.5% agarose gel was prepared with 1 ug of EDTA. The prepared particles were dialysed against 1 L water for 1 h. The final particles were mixed with a 6x gel loading dye and a total 20 ul sample was added to each well for all the different N/P ratios. The gel was run at 80 v for 30 min. The gel was imaged under UV trans-illuminator and photographs were taken using camera. The images were processed for clarity using Microsoft PowerPoint (Microsoft, USA).

### 2.4 Scanning Electron Microscopy (SEM)

The samples were prepared on coverslips. The coverslips were treated with 1M hydrochloric acid overnight and thoroughly rinsed with deionized water and air dried. The prepared particles were dropped onto the coverslips and air dried under fume hood. Part of the coverslips were for with a diamond cutter and foxed to a double sided carbon tape onto a metal stub. The samples were sputter coated with gold-palladium alloy for 40s (EMS sputter coater, USA). The samples were images under FESEM (Quanta650 ESEM, FEI, USA).

### 2.5 Cell culture

The MG63 cells (ATCC, USA) were cultured in low glucose down media (Gibco, USA) supplemented with 10%FBS (Gibco, USA), and 1% Penicillin-Streptomycin (Gibco, USA). The cells were treated with 0.25% Trypsin-EDTA (Gibco, USA) at 80% confluency. The cells were cultured in a BSL2 biosafety cabinet and incubated in a CO_2_ incubator (Cellexpert, Eppendorf, USA) at 5%CO_2_, 37 °C.

### 2.6 In vitro Transfection

Around 20,000 cells were seeded into 48 well plate (Celltreat, USA) and 20 μL of the sample equivalent to 100 ng of plasmid DNA was added to the cells and incubated for 24 h in a CO_2_ incubator at 5%CO_2_, 37 °C. The cells were imaged under fluorescence microscope (IX83, Olympus, USA) for brightfield and green fluorescence protein using with excitation at 488 nm.

### 2.7 In vitro toxicity

Around 10,000 cells were seeded into 96 well plate (Celltreat, USA) and the samples were added to the cells and incubated for 24 h in a CO_2_ incubator at 5%CO_2_, 37 °C. The media was removed and 10 μL (3-(4,5-dimethylthiazol-2-yl)-2,5-diphenyltetrazolium bromide (MTT) reagent with 100 μL media was added to each well and incubated for 2 h. The media was removed, and the formazan crystals were dissolved in 100 μL DMSO. The color change was read at 540 and background was read at 700 nm using a multi-mode plate reader (Biotek, USA).

### 2.8 Statistics

All experiments were carried out with triplicates. The data were shown as mean ± standard deviation. The significance of difference, *p<0.005 was evaluated doing One-way ANOVA (v5.03, GraphPad Prism, USA).

## 3. Results

The SMART process (Figure1A) is proposed for preparation of nanoparticles. The nanoparticles at different N/P ratios showed a particle size from 100 to 200 nm. The N/P 0 is the group with no pDNA but CB buffer. The N/P 1 is the formulation with higher pDNA which showed higher particle size owing to its loose binding of the Lipid-pDNA complexation. The groups N/P 5, 8, and 10 showed similar particle sizes of around 150 nm. Between these three groups, the N/P 8 formulation yielded smallest particle size of 138± 8nm (Figure 1B). The zeta potential analysis showed that all formulation except pDNA showed a positive value owing to the lipid complexation. The formulation N/P 1 showed a lower value compared to N/P 5, 8, and 10. The zeta potential value of 29 mV for N/P 8 (Figure 1C) indicates the stability of the formed particles.

**Figure 1.**
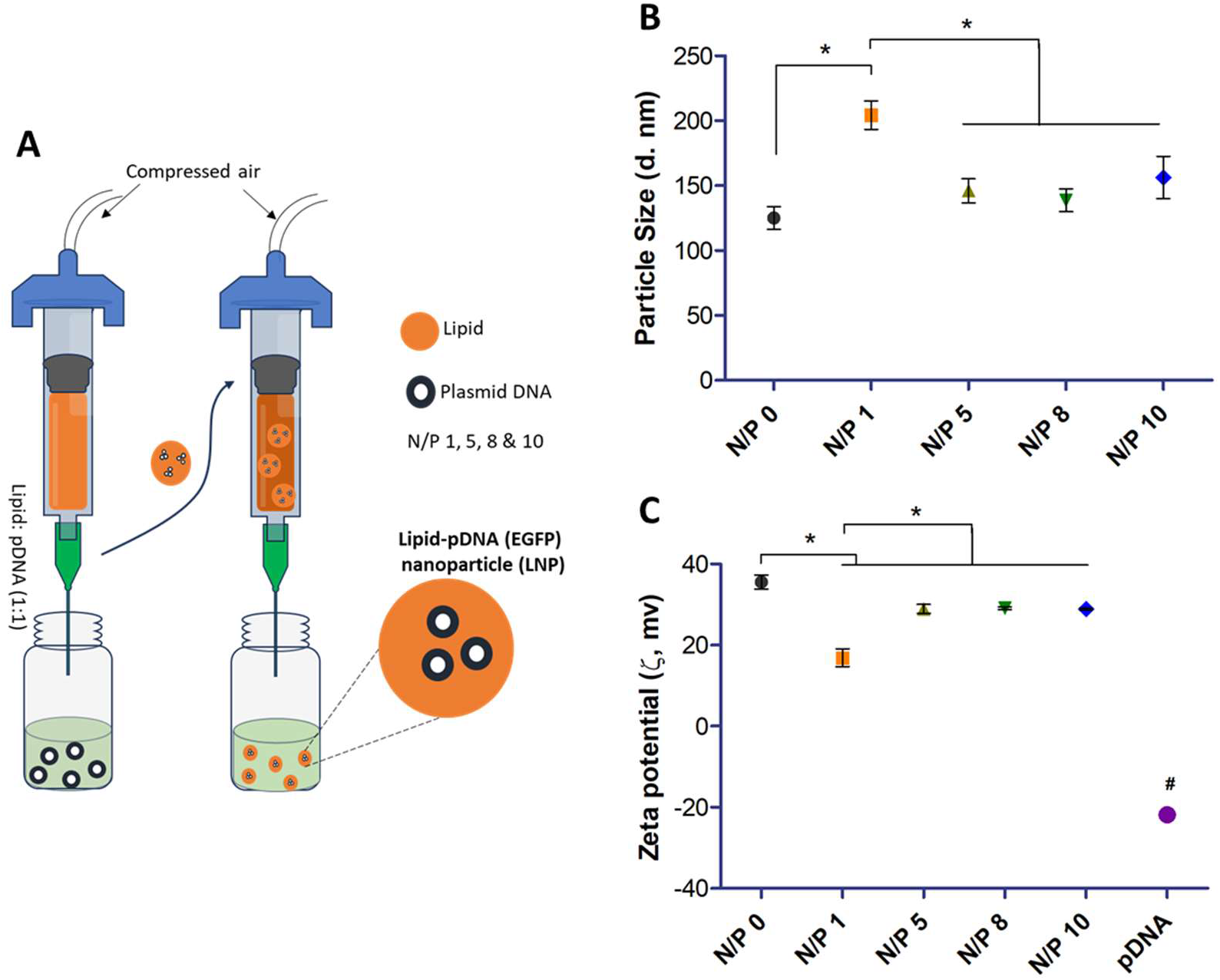
[A] Cartoon depicting the SMART process for preparation of lipid-pDNA (GFP) (LNP) particles with different N/P 1, 5, 8, & 10, [B] Particle size and [C] Zeta Potential. The data represented as mean ± standard deviation. The significance of difference between the groups was determined using One-way ANOVA with *p<0.005.

The SEM images (Figure 2A) showed the intact and smooth particles with clear shape and structure for all particles with N/P 1 to 10. The gel retardation assay (Figure 2B) showed no free plasmid DNA beyond N/P 1. The bright light in the wells shows that the plasmid DNA is not free to move down the gel signifies that all N/P ratios exhibited 100% encapsulation and complexation of pDNA. The difference in the brightness is due to the varying amounts of plasmid DNA to compensate the lipid component impeded to achieve the desired N/P ratios.

**Figure 2.**
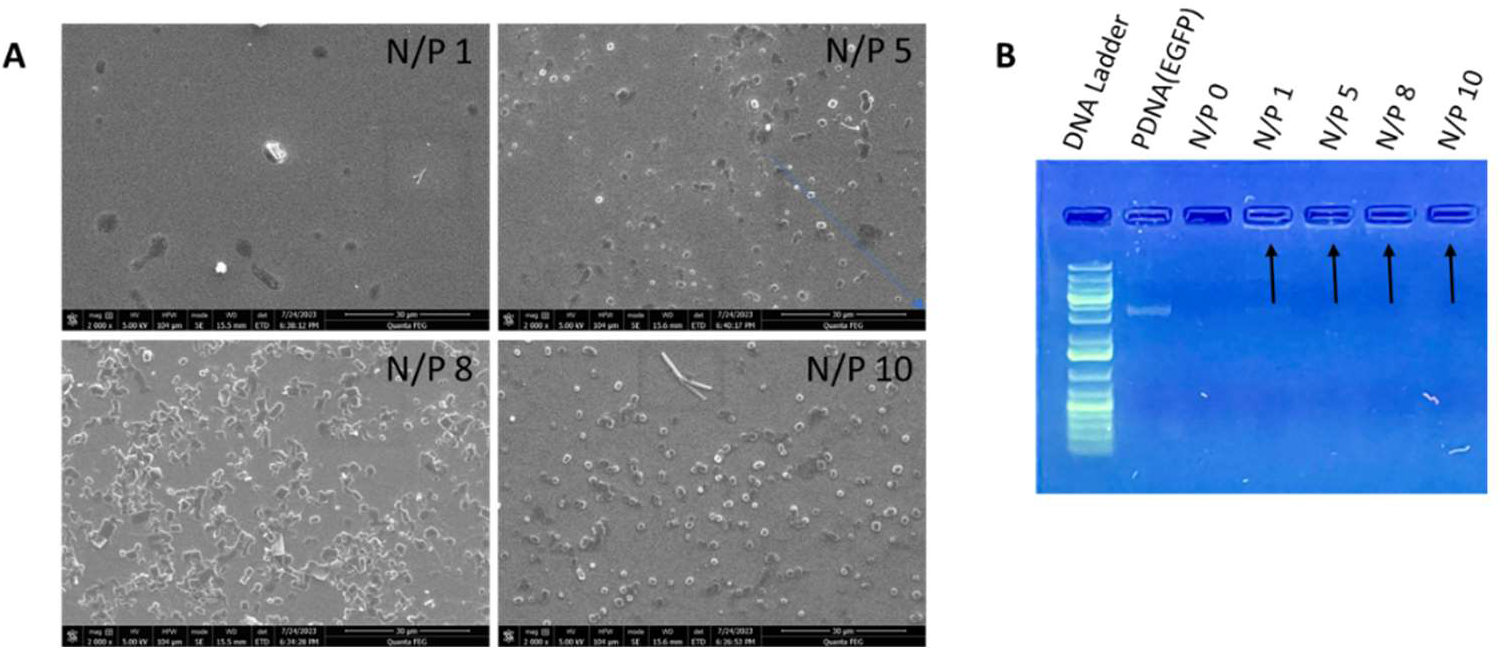
[A] SEM images of formulations of Lipid-pGFP at different N/P ratios and [B] Gel retardation assay showing no free pDNA was available with formulations revealing 100% encapsulation.

The MTT assay data in figure 3A shows the incremental toxicity with higher N/P ratio due to higher lipid component. Also, the varying amounts of the nanoparticles showed decrease in cell viability signifies the ability to tailor the dose amounts needed to elicit a therapeutic response. The cell viability is reduced to 60% overall compared to cells alone group could be attributed to higher amounts of lipids used for preparation of the particles.

**Figure 3.**
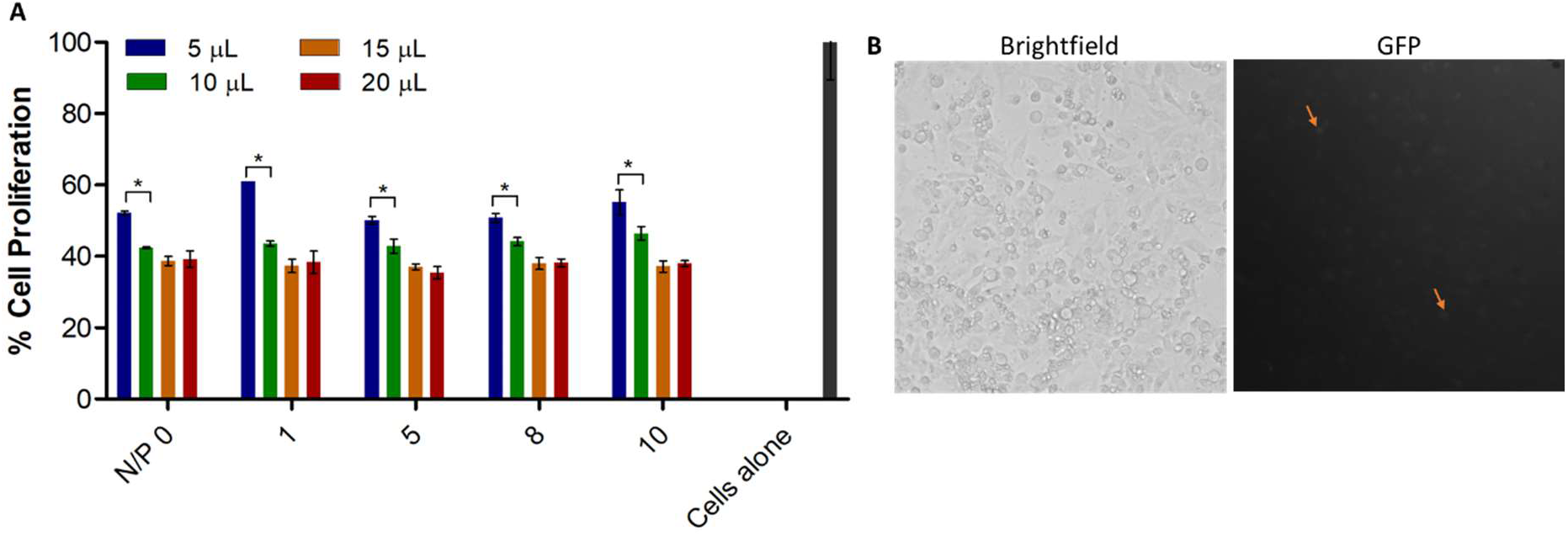
[A] Cell viability of Lipid-pDNA formulations in MG63 cells invitro and [B] Brightfield and GFP fluorescence (greyscale image) of N/P 8 at 20 μL showing expression of GFP (orange arrows). The data represented mean ± standard deviation. The significance of difference between the groups was determined using One-Way ANOVA with *p<0.005.

The fluorescence image (greyscale) (Figure 3B) shows the bright regions on the black background are the photons emitted by the expression of GFP inside MG63 cells (orange arrows). This result indicates that the uptake and internalization of the LNP occurred and the ionizable lipids facilitated the release of the pDNA from the particle without degradation into the cell and facilitated its transcription and translation.

## 4. Discussion

The SMART process is a portable and unique process that was employed at the minimal scale for this study to prove that the technology can fabricate the lipid-pDNA particles. The process achieved a particle size of around 138 nm with narrow size distribution for particular N/P formulation. Similar results of 131 nm were obtained with cationic lipid particles through thin film hydration method^5^. The lipid composition also plays a key role in particle size. The delivery vehicle performs to its optimum level owing to its composition, charge, and size. The lipid composition of DOTAP DSPC LNP – GFP yielded a particle size of 818 ± 161 nm showed a 94% encapsulation when fabricated with a microfluidic system process^25^. With similar composition, through SMART process we were able to produce the nanoparticles of 138 ± 8 nm.

The study also proved the ability of the SMART process to prepare an effective and functional gene delivery vehicle *in vitro*. The particles were stable based on the gene potential and showed 100% encapsulation and complexation achieving maximum encapsulation based on the gel retardation assay data. Ionizable lipid are crucial for electrostatic interaction of the lipid particle which are the sole interactions between the lipid and negatively charged pDNA. The addition of ionizable lipids is attributed for the 100% encapsulation of pDNA for all N/P formulations. The incorporation of ionizable lipid might have also compensated for the cell uptake leading to the transfection of GFP as occurred in the cell cytoplasm. The N/P 8 ratio showed optimal properties for a successful formulation for plasmid transfection. Ionizable lipids are known to be toxic due their ability of non-specific binding to proteins^26^. It was visible in the MTT assay for the lipid formulations in MG63 cells where all formulations showed around 60% reduce in viability with the least amount of formulation loading of 5 μL. More studies need to be carried out to fine-tune the lipid composition to minimize this cell death.

Overall, the results demonstrated successful particle preparation and functionality using SMART. We continue to explore the possibilities of this technology with more elaborate and robust experimental design for real-time applications in future.

## 5 Acknowledgement

The authors thank UT Austin and COP, UT Austin for central facilities. The authors thank Mei, Melissa, and Dr. Ghosh, COP, UT Austin for fluorescence microscopy facility.

